# Microscale dysfunction and mesoscale compensation in degenerating neuronal networks

**DOI:** 10.1101/2025.06.06.657602

**Authors:** Vegard Fiskum, Nicolai Winter-Hjelm, Nicholas Christiansen, Axel Sandvig, Ioanna Sandvig

## Abstract

Progressive neurodegenerative diseases involve neuronal dysfunction from cellular to circuit to whole-brain levels, but complexity and variability, both between and within diseases, pose significant research challenges. However, although they are differentiated by anatomical origins, vulnerable neuronal subtypes, and specific misfolded proteins, neurodegenerative diseases also share many important features. During presymptomatic disease phases, neural networks initiate multiple compensatory processes to maintain network function, including increased network centralisation and reliance on a rich-club of hub nodes, which have been proposed as common reconfigurations to neural network damage. Currently, while supporting evidence for such mechanisms has been found in some neurodegenerative diseases, it is limited in others, like ALS. This knowledge gap makes it challenging to ascertain if there are indeed common pre-symptomatic mechanisms within and across different neurodegenerative diseases. To address this, we investigated the structural and functional properties of ALS patient derived motor neuron networks and counterpart networks from a healthy donor using longitudinal multielectrode array recordings and graph theory-based network analysis. We demonstrate microscale-level motor neuron dysfunction, including TDP-43 proteinopathy, hyperactivity and reduced spike amplitude. Structurally, we observed neurite hypertrophy, indicating that degenerating networks attempt to establish new connections. We furthermore document mesoscale-level functional reconfigurations, including increased rich-club connectivity and network assortativity, indicating functional compensation in ALS where networks become more centralised to maintain computational capacity. We thus provide novel evidence that ALS networks become increasingly centralised, which places progressively mounting demands on a rich-club, predisposing networks to further damage, consistent with existing models of common reconfigurations in neurodegenerative disease.

**Significance statement:** This study makes significant contributions to preclinical modelling of neurodegenerative disease, with specific relevance for ALS research. By utilising human cellular models, longitudinal extracellular electrophysiology, and advanced network analysis, we show that known features of ALS can be recapitulated in *in vitro* engineered neural networks, and that these networks allow for novel hypothesis testing and identification of pre-symptomatic pathological processes including increased centralisation. This has previously been observed in other neurodegenerative diseases, but evidence has been limited in ALS. Our results advance our understanding of motor neuron network dynamics in ALS and contribute to a shared understanding of how neurodegenerative diseases affect neural networks which go beyond specific disease diagnosis, elucidating fundamental processes in neural network function and disease response.

## Introduction

Over the past decade, the role of brain network structure and function in progressive neurodegenerative disease has received increasing attention. Although different neurodegenerative diseases originate in different anatomical regions of the central nervous system and selectively affect certain vulnerable neuronal populations, there appear to be common patterns of changes in functional connectivity. These patterns include early compensatory changes in pre-symptomatic phases, which fail in increasingly detrimental ways as the diseases progress (Gregory et al. 2017). Even at the microscale, or cellular level, misfolded proteins and formation of aggregates are nearly ubiquitous features, although the specific proteins involved may differ, such as TAR DNA-binding protein 43 (TDP43) in amyotrophic lateral sclerosis (ALS) (Peng, Trojanowski, and Lee 2020). Understanding these shared responses may enable early detection, broad preventive strategies, and better treatments.

Neural networks organize their structural and functional features to facilitate high computational capacity while balancing metabolic demands and anatomical constraints, which can be characterized by graph theory (Bullmore and Sporns 2009, 2012). We have previously outlined that well-functioning neural systems achieve this balance by developing several hallmarks of efficient computation, including scale-freeness, small-world topology, and a modular organization (Heiney et al. 2021). This organisation facilitates local specialisation and global integration of information processing, the latter of which is mediated by a community of interconnected high degree hub nodes called a rich-club (Bullmore and Sporns 2012; Griffa and Van den Heuvel 2018). Maintaining these properties is vital for preserving network function, both at the mesoscale circuit level and at the macroscale whole-brain level, and disruptions can lead to loss of function, as seen in both traumatic brain injury and in neurodegenerative diseases.

Neural networks affected by progressive neurodegenerative disease initiate various processes to maintain network function, including changes in firing patterns (Pardillo-Diaz et al. 2022; Gunes et al. 2022; Bullmore and Sporns 2009) and structural and functional connectivity (Pievani et al. 2014), many of which precede clinical symptoms. A promising model outlined by Hillary et al. proposes that early structural degeneration results in increased activity and functional hyperconnectivity as a short-term adaptive response (Hillary and Grafman 2017; Hillary et al. 2015). In the longer term, these changes tend to make the networks more centralised, relying on a rich-club of hub nodes to facilitate a progressively larger fraction of information transmission (Hillary et al. 2014). While these changes may maintain function, they also place an increasing demand on a small subset of the network, which face increasing metabolic demands and become increasingly vulnerable to failure (Stam 2024). Alongside such functional network reconfigurations, networks may also initiate other degeneration-induced responses including enhanced neurite outgrowth (Saad, Segal, and Ayali 2015), which may contribute to maintaining network function for a time.

Self-organised *in vitro* neural networks mirror the behaviour of neurons and networks in the brain (Shein-Idelson, Ben-Jacob, and Hanein 2011; Poli, Pastore, and Massobrio 2015; Schroeter et al. 2015). Without behavioural and cognitive symptoms, graph theory can establish the network’s features consistent with high computational capacity (Heiney et al. 2021). By integrating these approaches with advanced neuroengineering and electrophysiology, we have also demonstrated that it is possible to probe micro- and mesoscale dynamic neural network reconfigurations caused by selectively induced perturbations (Weir et al. 2023), including neurodegenerative pathology (Valderhaug et al. 2020; Valderhaug et al. 2021).

In this study, we investigated the microscale and mesoscale dynamics of human ALS patient induced pluripotent stem cell (iPSC)-derived motor neurons (MNs) harbouring an endogenous expansion mutation in C9orf72 (ALS MNs) compared to healthy counterparts (HC MNs). We characterised the structural network qualities, assessed TDP-43 proteinopathy, and longitudinally assessed their functional connectivity by complex network analysis of extracellular electrophysiology using multielectrode arrays. We provide evidence of activity changes and, for the first-time, demonstrate that endogenous connectivity changes in ALS patient brain networks can be recapitulated in *in vitro* engineered neural networks, and that these compensatory properties drive the networks towards a more vulnerable state.

## Methods

### Experimental Design and Statistical Analysis

Human iPSCs were obtained from a healthy donor (female, 49 years, FA0000011 RUCDR Infinite Biologics and Target ALS) and a patient donor with confirmed ALS with *C9orf72* expansion mutation (female, 64 years, FA0000003 RUCDR Infinite Biologics and Target ALS), both reprogrammed according to the Sendai virus iPSC reprogramming method. iPSCs were cultured and differentiated into MNs according to the protocol by Nijssen *et al*. (Nijssen, Aguila, and Hedlund 2019), specifically the section “Procedure: E” regarding MNs from human iPSCs, with the exception that an orbital shaker was used throughout the embryoid body phase. The composition of cell media at different stages is shown in Table 1. Cells were seeded at day 10 following the start of the MN differentiation protocol. Each HD-MEA (n=4 for both HC and ALS) was seeded with 80 000 live cells, each well in the multiwell MEAs with 100 000 live cells (n=9 for HC and n=12 for ALS), wells in a 6-well plate were seeded with 800 000 MNs (n=4 for both HC and ALS), and each well in the 8-well chambered slides with 40 000 live cells.

**Table 1.** Cell media compositions during motor neuron differentiation. All cell media components were sourced according to Nijssen et al. (Nijssen, Aguila, and Hedlund 2019), and supplements were added to base media composed of the indicated mix of DMEM/F12-GlutaMax and Neurobasal.

|  | Days 1-2 | Days 3-9 | Day 10 | Days 11-12 | Day 13+ | Supplier | Product # |
| --- | --- | --- | --- | --- | --- | --- | --- |
| **DMEM/F12-GlutaMax** | 0,5 | 0,5 |  |  |  | Thermo Fisher | 31331028 |
| **Neurobasal** | 0,5 | 0,5 | 1 | 1 | 1 | Thermo Fisher | 21103049 |
| **N2** | 0.5X | 0.5X |  |  |  | Fisher Scientific | 12013479 |
| **B27** | 0.5X | 0.5X | 1X | 1X | 1X | Thermo Fisher | 17504044 |
| **Penicillin/Streptomycin** | 1X | 1X | 1X | 1X | 1X | Merck | P4333 |
| **Y-27632** | 5 μM | | 5 μM | | | Tocris | 6053 |
| **SB-431542** | 40 μM | | | | | Tocris | 1614 |
| **LDN-193189** | 200 nM |  |  |  |  | Tocris | 6053 |
| **CHIR-99021** | 3 μM | | | | | Tocris | 4423 |
| **Retinoic acid** |  | 200 nM | 200 nM |  |  | Sigma-Aldrich | R2625 |
| **SAG** |  | 500 nM |  |  |  | Tocris | 4366 |
| **Ascorbic acid** | 200 μM | 200 μM | 200 μM | 200 μM | 200 μM | Sigma-Aldrich | A4403 |
| **DAPT** | | | 10 μM | 10 μM | | Tocris | 2634 |
| **GDNF** |  |  | 10 ng/ml | 10 ng/ml | 10 ng/ml | Peprotech | 450-10 |
| **BDNF** |  |  | 10 ng/ml | 10 ng/ml | 10 ng/ml | Peprotech | 450-02 |

To compare the number of non-nuclear TDP-43 inclusions and network structural parameters in HC and ALS MN networks, we imaged immunolabelled TDP-43, heavy neurofilament and Hoechst nuclear staining in 20% of 8 separate networks from each group (n=720 images for both groups). Prior to quantification of TDP-43 inclusions we removed images with no identified nuclei or TDP-43 inclusions and normalized TDP-43 inclusion count to nuclei count for each image (final n=419 images for HC and n=463 images for ALS). Prior to quantification of network structural parameters, we removed images with no identified nuclei and normalized structural parameters to nuclei count for each image (final n=420 images for HC and n=558 images for ALS). Data point distributions of HC and ALS group levels of non-nuclear TDP-43 inclusions and network structure parameters were tested for normality by Kolmogorov-Smirnov test, and did not follow a normal distribution (Matlab R2024a, Mathworks). Group differences were therefore assessed by Wilcoxon’s Rank-sum test (Matlab R2024a, Mathworks), with Bonferroni correction for multiple comparisons.

Differences in HC and ALS MN network longitudinal electrophysiological activity was compared by generalised linear mixed models (GLMMs) using IBM SPSS Statistics (version 29.0.0.0). The models used genotype (HC or ALS) as a fixed effect with various network features as targets, utilising a linear model with repeated measurements of each network (the subjects of the model).

### Electrophysiological recordings

We studied MN networks over time both in high-density microelectrode arrays featuring 4096 recording electrodes (3Brain Arena, HD-MEAs), and 6-well MEAs with 64 recording electrodes each (Axion Biosystems M384-tMEA-6B, multiwell MEAs). For multiwell MEAs, network electrophysiological activity was recorded using an Axion Maestro Pro acquisition tool and AxIS Navigator software (version 3.12.2.2). Each recording lasted 30 minutes at 37°C and 5% CO2 with a sampling rate of 12.5 kHz. Prior to starting the recording, multiwell MEAs were left to settle for 15 minutes. The activity of all networks was recorded every other day from day 30 to day 40. For HD-MEAs, network electrophysiological activity was recorded using a Biocam Duplex System (3Brain) and BrainWave 5. Each recording lasted for 15 minutes at 37°C with a sampling rate of 18 857.72 Hz. Prior to starting the recording, HD-MEAs were left to settle for 5 minutes. The activity of all networks was recorded every other day from day 30 to day 46.

### Immunocytochemistry

Immunocytochemistry (ICC) was applied to confirm MN identity after differentiation and maturation of the iPSCs, assess network structure, and to investigate the presence of cytoplasmic inclusions of TDP-43. The approach was based on the work by Richter *et al*. (Richter et al. 2018). Cell media was removed, and the networks were fixed for 15 minutes in 3 % glyoxal solution consisting of 70.1% MQ H2O, 19.7% ethanol (Kemetyl 200-578-6), 7.8% glyoxal (40% wt. % in H2O, Sigma Aldrich 128465) and 0.75% acetic acid (Merck 1.00063). Then, the networks were washed with PBS (D8662, Merck) four times. The cells were then permeabilised with 0.5 % Triton-X (T8787, Merck) in PBS for 5 minutes. Networks were then washed three times with PBS, before adding a blocking solution of 5 % goat serum (PCN5000, Fisher Scientific) in PBS for 1 hour at room temperature on an orbital shaker at 30 rpm. After aspirating the blocking solution, primary antibodies in PBS with 5 % goat serum were added and left overnight on a shaker table at 4°C. The next day, primary antibodies were removed, and networks were washed four times with PBS. Secondary antibodies were then added in PBS with 5 % goat serum and left on an orbital shaker at 30 rpm for 3 hours, followed by nuclear staining for 10 minutes. The networks were then washed four times with PBS, before being washed once with milli-q water. The duration of each washing step was 5 minutes. All networks were fixed at 42 DIV.

MN networks were examined for expression of Islet-1 (ab109517, 1:250, Abcam), Chat (ab178850, 1:500, Abcam), HB9 (ab221884, 1:200, Abcam), NeuN (ab279295, 1:500, Abcam), Heavy neurofilament (ab4680, 1:1000, Abcam) and TDP-43 (PA5-27221, 1:500, Fisher Scientific), all visualized with the same secondary antibodies (Goat anti-mouse, ab150113, 1:500, Abcam; Goat anti-rabbit, ab175471, 1:500, Abcam; Goat anti-chicken, ab150171, 1:500, Abcam) and stained for Hoechst (62249, 1:2000, Fisher Scientific). All images were acquired using an EVOS M7000 microscope with the following light cubes: DAPI (AMEP4650), CY5 (AMEP4656), GFP (AMEP4651), and TxRed (AMEP4655) and the lens Olympus UPLSAP020x, 20x / 0.75 NA (N1480500).

### Image analysis

Five structural properties of MN networks, Neurite Branches, Junctions, Endpoints, Total and Average length, as well as identification of nuclei, were assessed using the NeuroConnectivity plugin for Fiji/ImageJ (Pani et al. 2014) using the settings in Table 2.

**Table 2.**
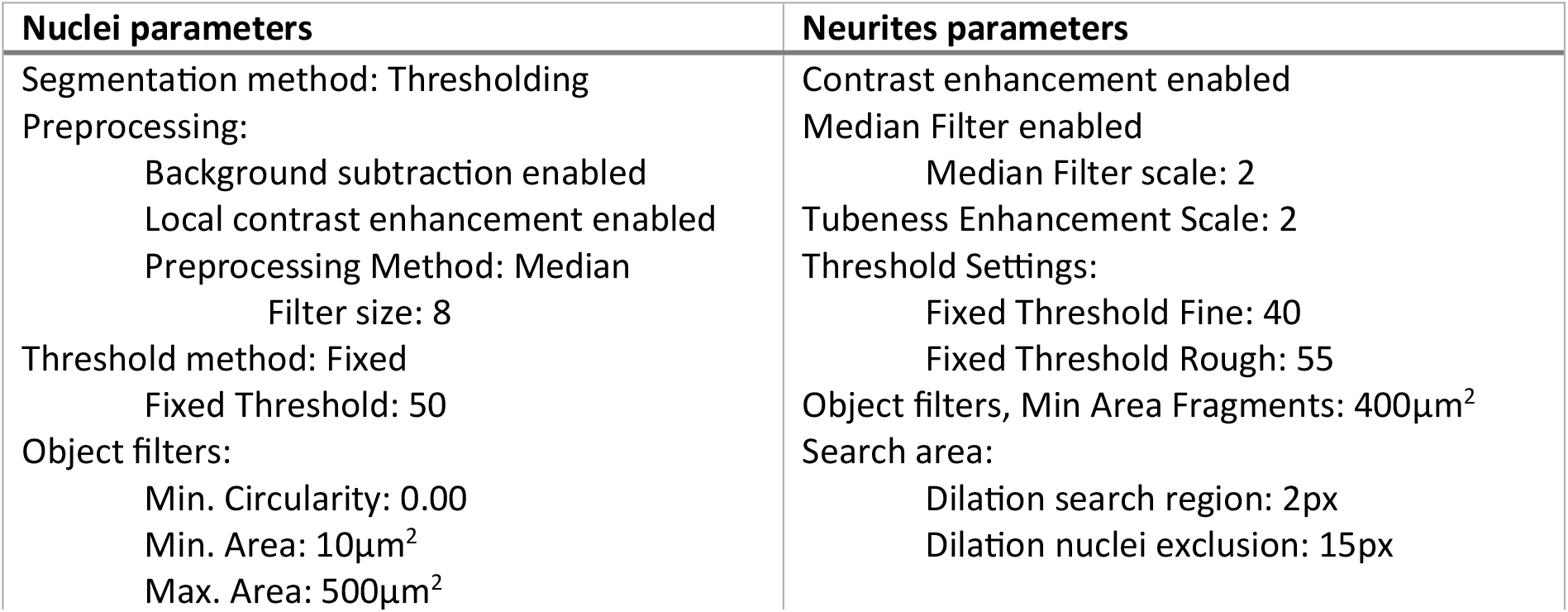
NeuroConnectivity parameters.

We used CellProfiler version 4.2.8 to identify TDP-43 inclusions not co-localised with nuclear labelling. First, Nuclei were identified using the Hoechst labelling by applying a Threshold, before identifying objects, using the settings in Table 3.

**Table 3.**
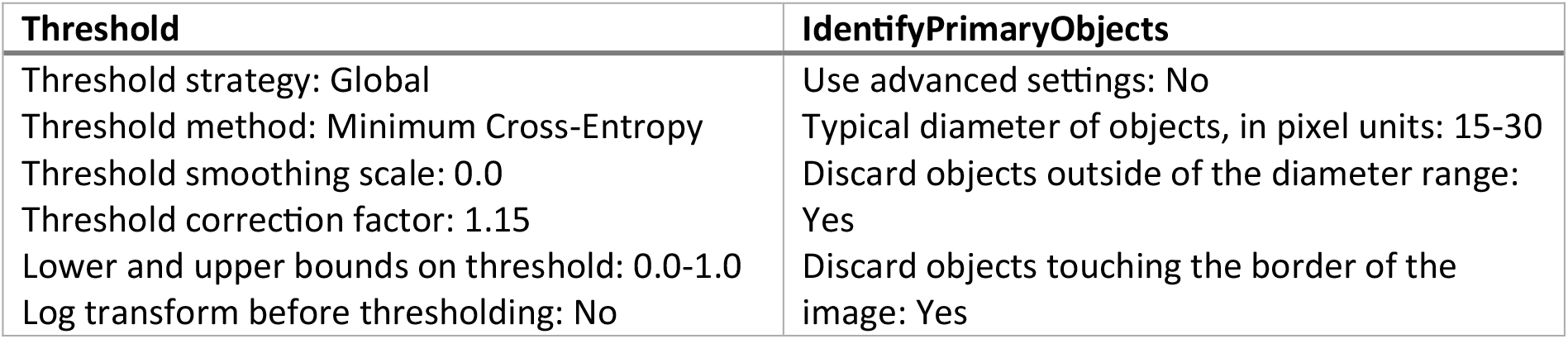
CellProfiler parameters for nuclei detection.

| Threshold | IdentifyPrimaryObjects |
| --- | --- |
| Threshold strategy: Global<br>Threshold method: Minimum Cross-Entropy<br>Threshold smoothing scale: 0.0<br>Threshold correction factor: 1.15<br>Lower and upper bounds on threshold: 0.0-1.0<br>Log transform before thresholding: No | Use advanced settings: No<br>Typical diameter of objects, in pixel units: 15-30<br>Discard objects outside of the diameter range: Yes<br>Discard objects touching the border of the image: Yes |

The threshold image was applied as an inverted mask to the TDP-43 labelling to isolate non-nuclear TDP-43 labelling, before applying a Threshold and identifying objects using the settings in Table 4.

**Table 4.** CellProfiler parameters for cytoplasmic TDP-43 inclusion detection.

| Threshold | IdentifyPrimaryObjects |
| --- | --- |
| Threshold strategy: Global | Use advanced settings: No |
| Threshold method: Otsu, Two classes | Typical diameter of objects, in pixel units: 10-20 |
| Threshold smoothing scale: 0.0 | Discard objects outside of the diameter range: Yes |
| Threshold correction factor: 2 | Discard objects touching the border of the image: No |
| Lower and upper bounds on threshold: 0.0-1.0 |  |
| Log transform before thresholding: No |  |

### Nuclear and cytoplasmic TDP-43 isolation

To identify the extent to which TDP-43 mislocalised from the nucleus to the cytoplasm of ALS patient derived MNs, we utilised a Thermo Scientific NE-PER Nuclear and Cytoplasmic Extraction Kit (Thermo Fisher Scientific 78833) with Thermo Scientific™ Halt™ Protease and Phosphatase Inhibitor Cocktail, EDTA-free (Fisher Scientific 10127963), following the manufacturer’s instructions. Briefly, cells from n=4 networks per group were harvested from 6-well plates with trypsin-EDTA (T4049, Merck) and centrifuged at 500g for 5 minutes. Cell pellets were washed by resuspension in PBS (D8662, Merck), before centrifugation at 500g for 3 minutes. The PBS was removed, before addition of protease and phosphatase inhibitor supplemented cytoplasmic extraction reagent 1. Cells were vortexed at the highest setting (3000rpm) for 15 seconds and incubated for 10 minutes before adding cytoplasmic extraction reagent 2. Cells were vortexed for 5 seconds before incubating for 1 minute, then centrifuged at 16 000g for 5 minutes. The supernatant containing the cytoplasmic extract was collected, before suspending the remaining cell pellet in protease and phosphatase inhibitor supplemented nuclear extraction reagent. The cells were vortexed for 15 seconds every 10 minutes for 40 minutes, before they were centrifuged at 16 000g for 10 minutes. The supernatant containing the nuclear extract was then collected. All centrifugations were performed at 4°C, and all samples, extracts, reagents, vials and pipette tips were kept on ice throughout the process. Extracts were stored at -80°C until they were analysed by western blot, using an antibody for TDP-43 (PA5-27221, 1:1000, Fisher Scientific), and normalising protein levels against actin levels. The fraction of cytoplasmic to nuclear TDP-43 was then compared between HC and ALS MN networks.

### Data analysis

For multiwell MEAs, raw data was filtered using a Butterworth high-pass filter of 200 Hz and a Butterworth low-pass filter of 3 kHz. Spikes were identified using adaptive threshold crossing with a 6 standard deviation threshold, with pre-spike duration of 0.84 ms and post-spike duration of 2.16 ms, coincidence occurrence threshold of 4 electrodes and coincidence event window of 80 µs. Wells with a network firing rate less than 5 spikes per minute or with fewer than 7 active electrodes were excluded from further analysis, resulting in n=9 networks for HC and n=12 networks for ALS. The median firing rate was determined by the number of spikes recorded for a single electrode divided by the total recording time, then taking the median of each network. To generate network connectivity matrices of multiwell MEA networks, the Pearson correlation of electrode pairs was used to generate a weighted graph, before removing 90% of the weakest connections to prune trivial connections before calculation of network features.

For HD-MEAs, n=4 for both healthy and ALS conditions, raw data was filtered using a 5^th^ order Butterworth high-pass filter, removing low frequency noise (below 200 Hz). Spike detection was conducted using the PTSD algorithm (Maccione et al. 2009). The threshold was set to 8 times the standard deviation of the noise, the peak lifetime period duration to 1.5 ms and the refractory time to 1 ms. Filtering and spike detection was performed in BrainWave 5 (3Brain). The median firing rate was determined by the number of spikes recorded for a single electrode divided by the total recording time, then taking the median of all electrodes for each network. To generate the network connectivity matrix of the HD-MEA recordings, spike times were separated into 100 ms bins. Then, the co-occurrences of the binned spike times were identified, resulting in a count of the number of times spikes on different electrodes co-occur. Stronger connections will have a higher tendency to co-occur. To establish a threshold for non-spurious connectivity, randomized series of corresponding data were generated by shuffling the original data. This was repeated 10 times, and co-occurrence counts which were equal to or lower than the mean of the shuffled data was removed from the connectivity matrix. Of these connections, the 1 % of strongest connections were further selected, resulting in binary connectivity matrixes with a comparable number of connections. Finally, the giant component from this was analysed using common network metrics.

Small-world propensity provides an estimate to how well each network conforms to small-world principles (I.e. high clustering and low average path length), as described in (Muldoon, Bridgeford, and Bassett 2016). A network with small-world propensity above 0.6 is considered small-world. The mean rich-club coefficient measures the tendency for nodes with high degree to interconnect (McAuley, da Fontoura Costa, and Caetano 2007). Assortativity is a measure of the degree correlation of connected network nodes, and network density is the proportion of possible edges which are present in the network. Community detection was done using CDlib (Rossetti, Milli, and Cazabet 2019), while modularity evaluation was done using NetworkX (Swart 2008) according to (Clauset, Newman, and Moore 2004). The Matlab function fit, with fitType=‘power1’ was used to assess if node degree distribution followed a power law (Matlab 2020b, MathWorks).

## Results

### ALS motor neurons exhibit endogenous TDP-43 proteinopathy

ICC of HC and ALS MN networks confirmed expression of MN markers Islet1, HB9 and ChAT alongside pan-neuronal marker NeuN, shown in Figure 1.

**Figure 1.**
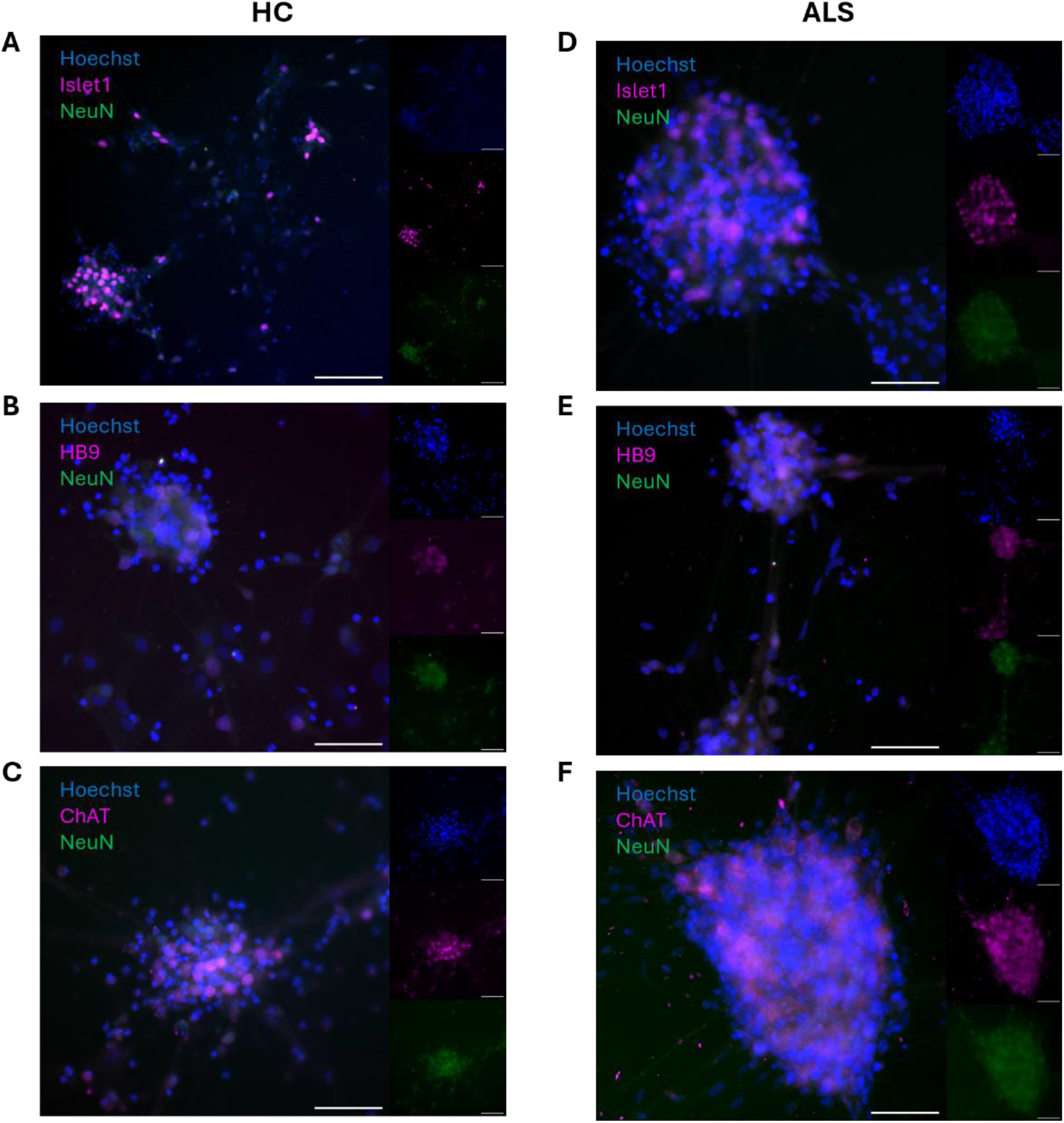
Immunocytochemistry confirms motor neuron cell identity. Both healthy control (HC) and ALS patient derived motor neuron networks showed expression of motor neuron specific markers Islet1 (A and D), HB9 (B and E) and ChAT (C and F) alongside pan-neuronal marker NeuN. Scale bar = 100 µm.

To assess if ALS MN networks showed signs of TDP-43 mislocalisation and proteinopathy, we performed separation of cytoplasmic and nuclear cell fractions to assess the levels of TDP-43 protein in subcellular locations in HC and ALS MN networks by western blot. As shown in Figure 2A, we found a 7.415-fold increase in the fraction of cytoplasmic to nuclear TDP-43 in ALS MN networks compared to HC MN networks. Furthermore, ICC-labelling of cytoplasmic TDP-43 positive protein inclusions, as shown in Figure 2B, showed a significant increase in the number of aggregates in ALS MN networks compared to HC MN networks (z-value=-9.6858, p= 3.47×10^−22^).

**Figure 2.**
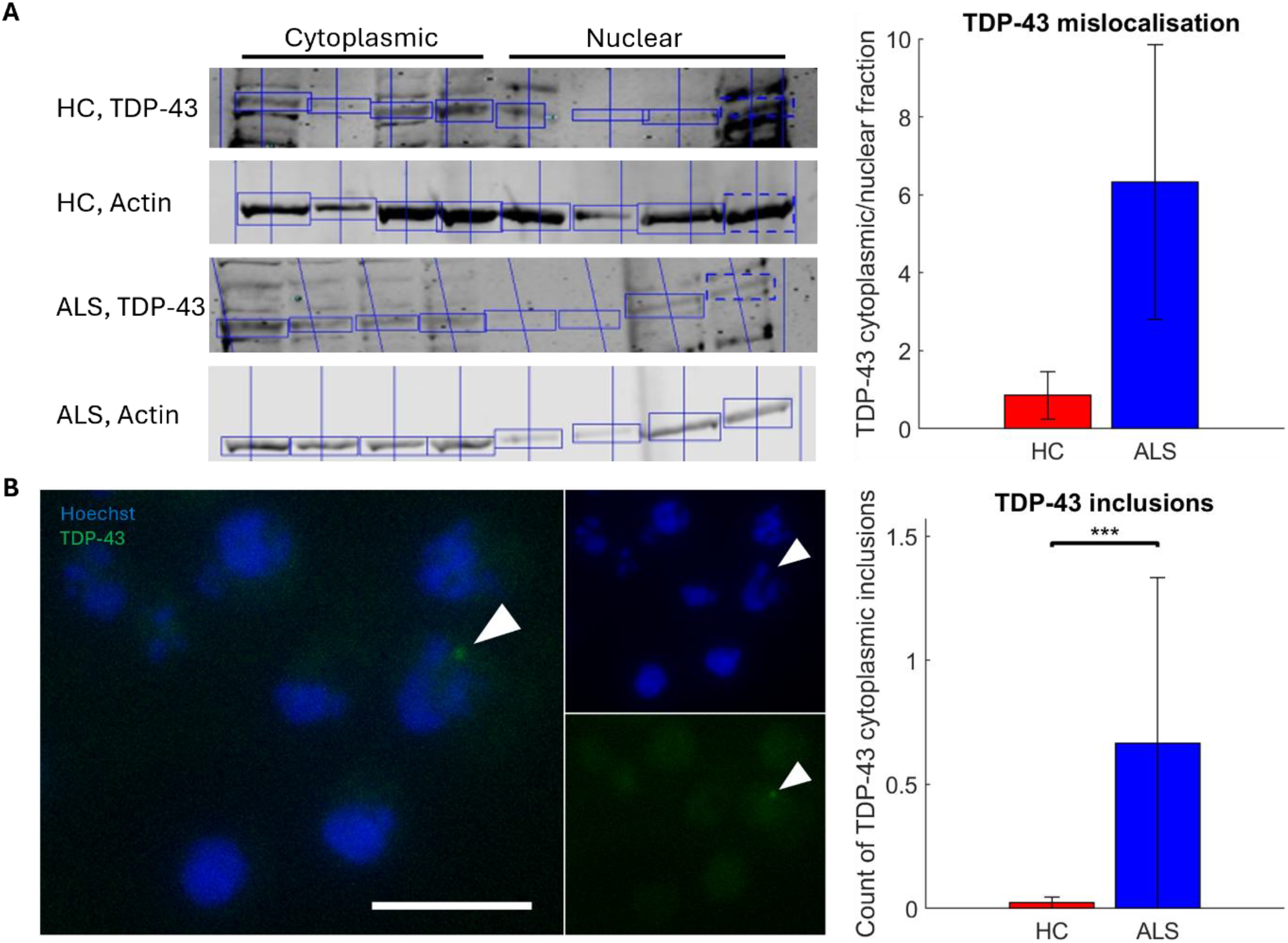
ALS motor neuron networks exhibit TDP-43 proteinopathy. Western blot analysis of levels of TDP-43 showed that ALS MN networks had a 7.415-fold increase in the cytoplasmic to nuclear protein ratio compared to HC counterparts, n=4 for HC and ALS (A). White arrows highlight inclusions of TDP-43 (green) accumulating outside of the nucleus (blue) in ALS motor neurons (B). Scale bar = 25µm. The number of non-nuclear TDP-43 inclusions was significantly increased in ALS MN networks compared to HC motor neuron networks, normalised to nucleus count (p=3.47×10-22), HC n=419, ALS n=463. *: p<0.05, **: p<0.01, *** p<0.001.

### ALS motor neuron networks exhibit microscale dysfunction and mesoscale compensation

MN network activity from HD-MEAs and multiwell MEAs is presented in Figure 3 and Figure 4 with longitudinal medians and median absolute deviations of each measurement to the left, and GLMM estimated group averages with 95% confidence intervals (CI) with statistical comparison to the right.

**Figure 3.**
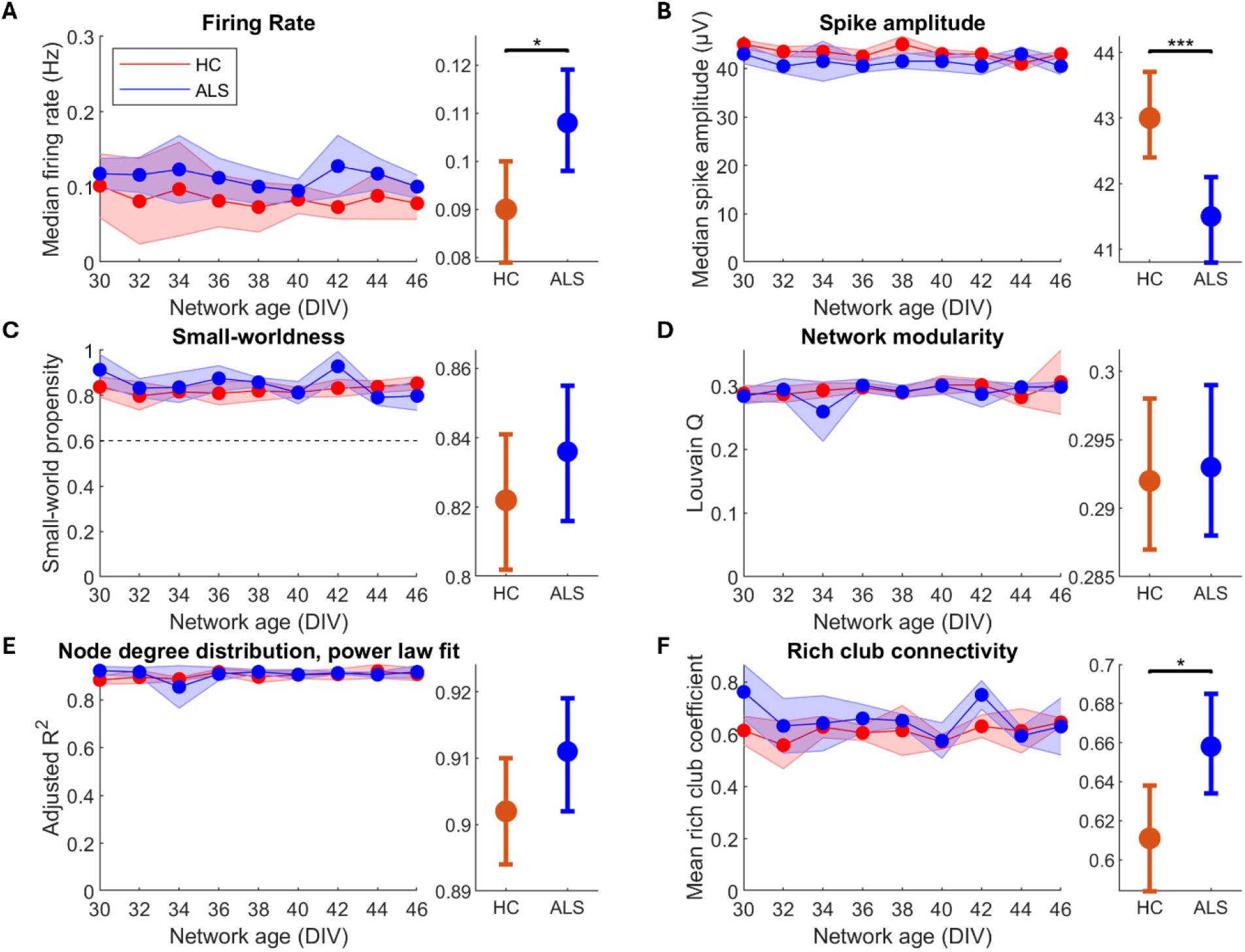
ALS patient derived and healthy motor neurons on HD-MEAs self-organise into networks with features of high computational capacity. 9 HD-MEA recordings per network (n=4 for both HC and ALS) in the period 30-46DIV, A-F left, were compared between HC and ALS groups using a generalised linear mixed model (GLMM), A-F right. Compared to HC, ALS MN networks had higher firing rate (A, p= 0.0140), lower spike amplitude (B, p= 1.09×10-4), similar small-world propensity (C, p= 0.315), similar modularity (D, p= 0.831), similar degree distributions which follow a power law, indicating scale-free networks (E, p=4.11×10-3), and increased mean rich-club coefficient (F, p=0.0180). The similarity of the features shown in C-E indicate that healthy and ALS MN networks both self-organised into networks with high computational capacity. Left plots show longitudinal medians with shaded regions indicating median absolute deviation. Right plots show GLMM estimated group averages with 95% confidence intervals. *: p≤0.05, **: p<0.01, ***: p<0.001.

**Figure 4.**
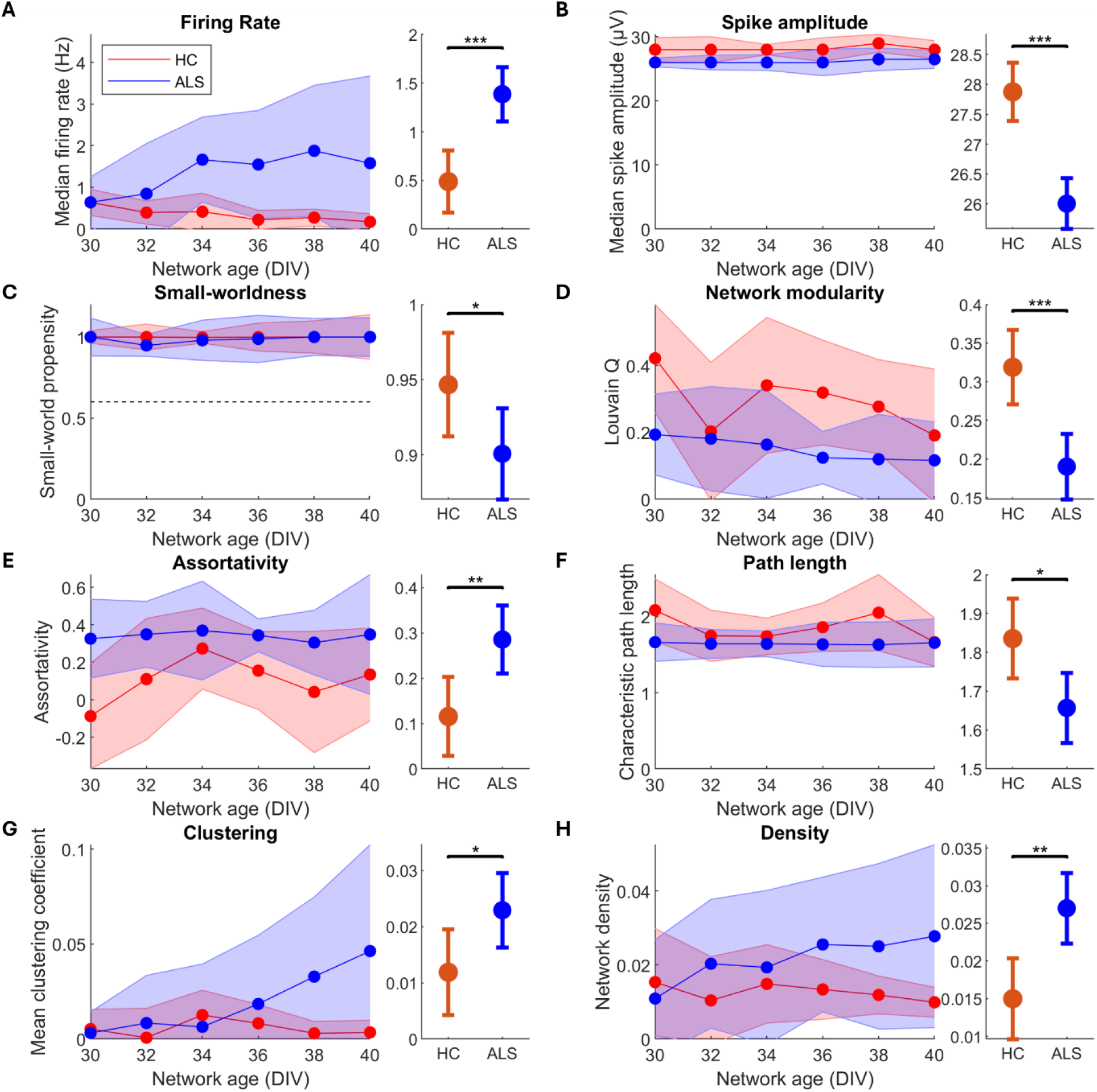
ALS patient derived motor neurons self-organise into networks with significant deviations from healthy counterparts on multiwell MEAs. 6 multiwell MEA recordings per network (n=9 for HC and n=12 for ALS) in the period 30-40 DIV, A-H left, were compared between HC and ALS groups using a generalised linear mixed model (GLMM), A-H right. Compared to HC, ALS MN networks had higher firing rate (A, p=5.01×10-5), lower spike amplitude (B, p=8.07×10-4), lower small-world propensity (C, p=0.0498), decreased modularity (D, p=1.21×10-4), higher assortativity (E, p=4.11×10-3), shorter characteristic path length (F, p=0.0111), increased clustering (G, p=0.0324), and increased density (H, p=1.07×10-3). Left plots show longitudinal medians with shaded regions indicating median absolute deviation. Right plots show GLMM estimated group averages with 95% confidence intervals. *: p≤0.05, **: p<0.01, ***: p<0.001.

MN network activity from HD-MEAs was recorded every other day from 30 to 46 DIV, for a total of 9 repeated measurements per network, shown in Figure 3. GLMM results of activity from HD-MEAs are summarised in Table 5. The median firing rate was significantly higher in ALS MN networks compared to HC MN networks, as seen in Figure 3A. Simultaneously, ALS MN networks exhibited significantly lower spike amplitudes compared to HC MN networks, shown in Figure 3B. Both HC and ALS networks showed spontaneously emergent properties associated with highly computationally competent networks. This included similar levels of small-worldness, as seen in Figure 3C, where all networks had small-world propensity > 0.6 and similar degrees of modularity, shown in Figure 3D. The node degree distributions of the networks adhered well to a power-law fitting, as seen in Figure 3E, which indicates that both HC and ALS MN networks were scale-free. Power law exponents were not significantly different between HC and ALS MN networks (p=0.281, data not shown). Individual degree distributions are shown in Supplemental Figure 1. These traits are consistent with high computational capacity as outlined in (Heiney et al. 2021). Lastly, it is noteworthy that the ALS MN networks had a significantly higher rich-club coefficient compared to the HC MN networks, shown in Figure 3F. These features indicate mesoscale compensation involving increased dependency on a subset of highly interconnected nodes to maintain network computational capacity.

**Table 5.**
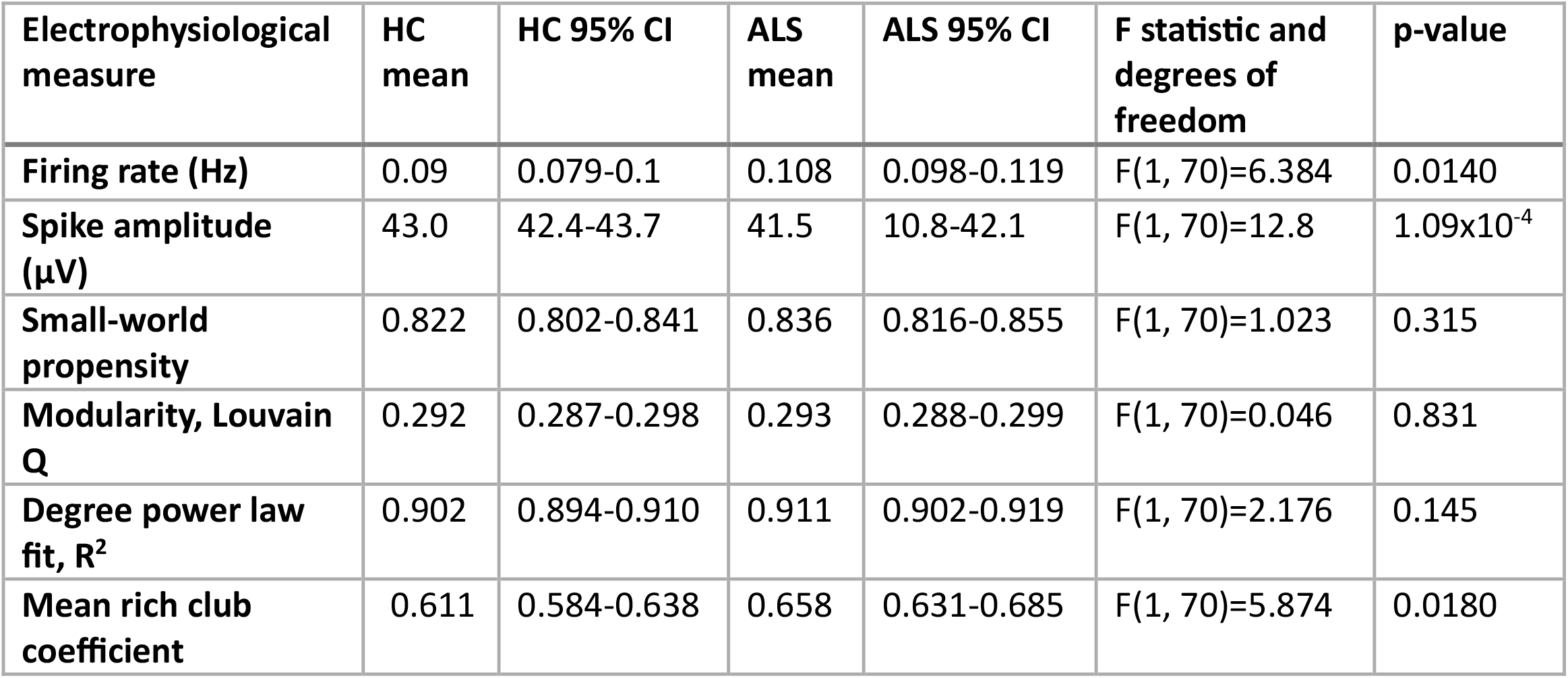
GLMM estimates and test results, HD-MEAs.

MN network activity from multiwell MEAs was recorded every other day from 30 to 40 DIV, for a total of 6 repeated measurements per network, shown in Figure 4. GLMM results from multiwell MEAs are summarised in Table 6. Similar to HD-MEA results, ALS MN networks had significantly higher median firing rate than HC MN networks, demonstrating network hyperactivity, and significantly lower spike amplitude, shown in Figure 4B. Furthermore, the ALS MN networks had significantly reduced small-world propensity and modularity compared to HC MN networks, shown in Figure 4C and D, indicating a disease-associated deterioration in computational capacity. Furthermore, we found that ALS MN networks had significantly higher assortativity than HC MN networks, as well as significantly decreased path length and significantly increased clustering shown in Figure 4E, F and G. Lastly, ALS MN networks had significantly higher density compared to HC MN networks, shown in Figure 4H.

**Table 6.** GLMM estimates and test results, multiwell MEAs.

| Electrophysiological measure | HC mean | HC 95% CI | ALS mean | ALS 95% CI | F statistic and degrees of freedom | p-value |
| --- | --- | --- | --- | --- | --- | --- |
| Firing rate (Hz) | 0,488 | 0,169-0,807 | 1,386 | 1,109-1,664 | F(1, 121)=17,689 | 5,01x10<sup>-5</sup> |
| Spike amplitude (μV) | 27,9 | 27,4-28,4 | 26,0 | 25,6-26,4 | F(1, 121)=32,651 | 8,07x10<sup>-8</sup> |
| Small-world propensity | 0,947 | 0,912-0,981 | 0,901 | 0,870-0,931 | F(1, 120)=3,926 | 0,498 |
| Modularity, Louvain Q | 0,319 | 0,471-0,367 | 0,190 | 0,148-0,232 | F(1, 120)=15,783 | 1,22x10<sup>-4</sup> |
| Assortativity | 0,116 | 0,0286-0,203 | 0,286 | 0,210-0,361 | F(1, 112)=8,585 | 4,11x10<sup>-3</sup> |
| Characteristic path length | 1,84 | 1,73-1,94 | 1,66 | 1,57-1,75 | F(1, 120)=6,660 | 0,0111 |
| Mean clustering coefficient | 0,0119 | 0,00424-0,0195 | 0,0229 | 0,0163-0,0296 | F(1, 120)=4,687 | 0,0324 |
| Network density | 0,0150 | 0,00964-0,0203 | 0,0270 | 0,0223-0,0317 | F(1, 120)=11,245 | 1,07x10<sup>-3</sup> |

### ALS motor neuron networks exhibit increased outgrowth of shorter neurites

We assessed structural features of MN networks by immunolabelled heavy neurofilament and Hoechst-labelled nuclei to determine if ALS MN networks deviated from HC MN networks. As shown in Figure 5C, ALS MN networks had significantly more neurite branches (z-value=-4.977, p=6.45×10^−7^) and junctions (z-value=-5.589, p= 2.29×10^−8^), as well as significantly higher total neurite length (z-value=-3.884, p= 1.03×10^−4^), while the average neurite length was significantly shorter (z-value=7.358, p= 1.87×10^−13^) compared to HC MN networks. The number of neurite endpoints was not significantly different between the two groups (z-value=-2.256, p= 0.0241).

**Figure 5.**
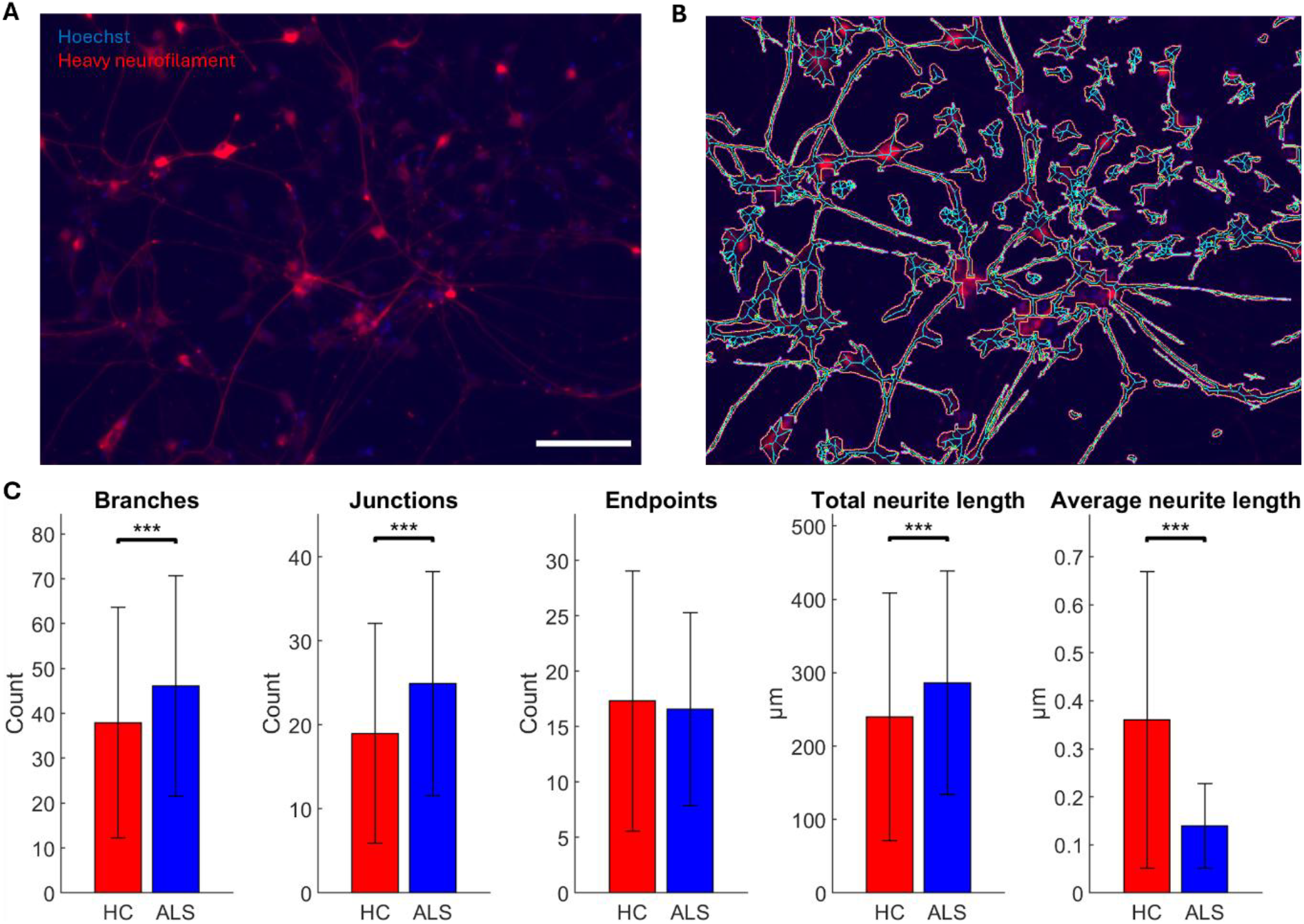
ALS motor neuron networks have extensive structural deficits compared to healthy counterpart. Motor neuron (MN) network structure was assessed by immunocytochemistry labelling of heavy neurofilament and nuclear Hoechst staining (A), using the NeuroConnectivity plugin for Fiji/ImageJ (Pani et al. 2014) to automatically identify neurites and nuclei (B). Quantification of structural features, normalised by nuclei count (C), showed that ALS MN networks had significantly more neurite branches (p=6.45×10-7) and junctions (p=2.29×10-8) than HC MN networks. At the same time, ALS MN networks had significantly higher total neurite length (p=1.03×10-4), while the average neurite length was significantly shorter (p=1.87×10-13) than in HC MN networks. The number of neurite endpoints was not significantly different (p=0.0241). HC n=420, ALS n=558. *: p<0.01, **: p<0.002, *** p<0.0002. Scale bar = 100 µm.

## Discussion

In this study we demonstrate for the first time that iPSC derived neural networks from a patient with confirmed neurodegenerative disease show endogenous pathological changes in functional connectivity. Across both HD-MEAs and multiwell MEAs, ALS MNs demonstrated microscale dysfunction, involving both hyperactivity, shown in Figure 3A and Figure 4A, and decreased spike amplitude, shown in Figure 3B and Figure 4B, compared to HC MNs. Neuronal hyperexcitability is a widely reported feature of ALS (Wainger et al. 2014; Ronchi et al. 2021; Sommer et al. 2022) and other neurodegenerative diseases (Pardillo-Diaz et al. 2022; Chen, Chehade, and Chu 2025; Targa Dias Anastacio, Matosin, and Ooi 2022). Furthermore, the significantly lower spike amplitude in ALS MN networks, shown in Figure 3B and Figure 4B, is consistent with previous observations in ALS patient-derived MN networks (Sommer et al. 2022). Lower spike amplitude reduces Ca^2+^ influx (Scarnati et al. 2020) and may represent an adaptation of ALS MN networks to minimise Ca^2+^-induced excitotoxicity and oxidative stress. However, this also reduces the amount of neurotransmitter released (Scarnati et al. 2020) and may contribute to disruption of network function. Overall, at the microscale, we found that ALS MN networks generate more, but weaker spikes, consistent with higher metabolic cost and increased oxidative stress.

Complex network analysis showed that ALS MN networks had multiple deviations from HC networks which indicate mesoscale compensation, including increased Rich-club coefficient, shown in Figure 3F, and Assortativity, shown in Figure 4E. This suggests that ALS MN networks become more centralised, and especially interconnected amongst hub nodes, resulting in more vulnerable networks. When local connections are impaired, either by loss of nodes or loss of signal fidelity as seen in the reduced spike amplitude in our findings, networks adapt by reorganising their functional connectivity. Because high degree hub nodes have high connectivity and activity, they are probabilistically more available for these adaptations, which therefore leads to an increased amount of information routing through hubs (Hillary et al. 2014; Roy et al. 2017). In an early phase of this adaptation, or if the network damage is acute and requires finite reconfiguration, this may be beneficial for maintaining network communication (Hillary and Grafman 2017). However, such reconfiguration increases the metabolic demand placed on hub nodes, and ongoing network damage such as in neurodegenerative disease leads to a continuous mounting stress on these nodes, which already represent some of the most metabolically costly components of the network (Bullmore and Sporns 2012; van den Heuvel et al. 2012). Eventually, the nodes succumb to this burden, and since so much of the network’s overall communication is dependent on them, the network fails to maintain its function without them (Stam 2014). While supporting evidence for such a pattern of damage and compensation has been found in cases of traumatic brain injury and Alzheimer’s disease (Hillary et al. 2015), evidence for this has been limited in ALS pathology until now. Previous work by Sorrentino et al. has shown that ALS patient brain networks become more centralised as the disease progresses (Sorrentino et al. 2018). We expand on this by showing increased rich-club connectivity in ALS MN networks on HD-MEAs. Our findings of increased network assortativity in multiwell MEAs furthermore indicate a shift towards a stricter hierarchy of node degree within the ALS MN networks. This is in line with both EEG (Iyer et al. 2015) and fMRI (Fekete et al. 2013) results from ALS patients.

Networks with greater assortativity are more likely to have a highly interconnected rich-club (Rubinov and Sporns 2010), thus supporting our findings in HD-MEA networks and providing clear evidence of pathological dynamics of centralisation in ALS MN networks.

Furthermore, we found increased network density in ALS, indicating a compensatory response to maintain network function by establishing new connections. While we did find significant decreases in ALS MN network small-worldness and modularity in multiwell MEAs, shown in Figure 4C-D, these findings were not replicated in HD-MEAs, shown in Figure 3C-D. Also, we did not observe any differences in degree distributions on HD-MEA networks, shown in Figure 3E. Together, these differences from HC MN networks, and lack thereof, indicate that the ALS MN networks were in a state of compensatory reconfiguration, where adaptive mechanisms appeared to maintain network function, with only minor detrimental effects of the underlying pathology. Our results thus provide novel findings of early functional network changes in ALS.

We demonstrated clear signs of TDP-43 proteinopathy in ALS MN networks by two separate methods. Following separation of nuclear and cytosolic cell fractions, western blot analysis found a 7.415-fold increase in the fraction of cytoplasmic to nuclear TDP-43, shown in Figure 2A. ICC labelling of TDP-43 also found a significantly higher number of non-nuclear TDP-43 inclusions, shown in Figure 2B. Cytoplasmic inclusions of TDP-43 are one of the most ubiquitous signs of MN cytopathology in ALS and frontotemporal dementia (Neumann et al. 2006; Arai et al. 2006), and tend to be aggravated by increased oxidative stress, reduced antioxidant capacity, and mitochondrial dysfunction (Kara et al. 2023). There is a substantial body of evidence supporting that MNs affected by ALS operate in a state of elevated oxidative stress (Cunha-Oliveira et al. 2020), and that environmental factors which further increase oxidative stress levels can contribute to risk and aggravation of the pathology (D’Amico et al. 2013). Combined, these findings clearly demonstrate TDP-43 proteinopathy and signs of cellular pathology, indicating that ALS MN networks operate at a higher level of oxidative stress and metabolic cost.

Structural analysis found that ALS MNs exhibited increased neurite outgrowth, but proportionally diminished long-range connections, shown in Figure 5C. These changes support our functional findings, indicating ongoing network reconfigurations in the face of neurodegenerative pathology. Indeed, the structural changes may offer an explanation to our multiwell MEA findings of decreased path length and increased clustering, shown in Figure 4F-G, despite the decrease in small-worldness, shown in Figure 4C. Since small-worldness is associated with low path length and high clustering (Muldoon, Bridgeford, and Bassett 2016), these findings seem contradictory. However, given our structural findings of increasing neurite density and shorter overall neurite length, alongside our findings of increased functional density, it is possible that a sizable number of short-range connections skew the group average path length and clustering. Notably, small-world propensity, as opposed to small-world index, accounts for properties such as density (Muldoon, Bridgeford, and Bassett 2016), allowing us to capture such seemingly counter-intuitive network dynamics. Increasing functional connectivity, at least in part caused by increased spiking activity, as a compensatory response to structural network damage has been observed both in acute injury and chronic disease and can result in higher clustering and shorter path length (Aswendt and Hoehn 2023). However, findings indicate that hyperactivity, hyperconnectivity, and proteinopathy are reciprocally linked, and that the compensatory responses described above can result in mounting metabolic demand, oxidative stress, hub overload and ultimate network failure (Hillary and Grafman 2017).

Here, we studied ALS MNs with an endogenous expansion in *C9orf72*, i.e., distinct from CRISPR-induced genetic predisposition or induced pathology by means of misfolded protein seeds. Future studies should examine MN networks derived from ALS patients with various genetic predispositions, as well as from patients with sporadic ALS, and asymptomatic donors with ALS-associated mutations, to assess if our findings are general trends of ALS or specific to certain patient groups. It is also possible that patients with the same identified genetic predisposition may not share the same network features due to other factors including epigenetic imprints. In this case, iPSC-based network models in combination with complex network analysis, and with network models developed by cell reprogramming that preserves epigenetic signatures by bypassing the stem cell stage, can contribute to personalised medical assessment (Okano and Morimoto 2022). Lastly, the methods and principles utilised in this paper can be applied to model and assess network dysfunction in other neurodegenerative diseases and to uncover which features of neural network dynamics, if any, may be shared by multiple pathologies.

In conclusion, in this study we show, for the first time, that *in vitro* neural networks derived from a patient with confirmed neurodegenerative disease exhibit endogenous functional network changes which reflect changes observed in multiple pathological conditions *in vivo*. We provide clear evidence of microscale dysfunction and mesoscale compensation in ALS MN networks, including new evidence of increased network centralisation. ALS networks also exhibit elevated cytoplasmic TDP-43 protein fraction and higher levels of cytoplasmic TDP-43 inclusions, showing that common pathological hallmarks of ALS are present within such networks. At the microscale, ALS networks exhibited hyperactivity despite weaker, less robust signalling. At the mesoscale level, the networks showed increased assortativity and connectivity within a rich-club, indicating increased centralisation in the face of ongoing network damage, which has been shown in other neurodegenerative diseases but has not previously been clearly demonstrated in ALS. Taken together, our findings provide novel evidence that ALS MN networks exhibit classical ALS cytopathology and self-organize into computational competent networks, but with compensatory hyperactivity and increased centralisation.

## Acknowledgements

This work was supported by the Olav Thon Foundation, ALS Norge and Alf Harborg’s fund. Western blot analysis of nuclear and cytoplasmic cell fractions was done by Proteomics and Modomics Experimental Core, PROMEC, at NTNU and the Central Norway Regional Health Authority.

## Author contributions

VF formulated the research questions and planned the experiments and methodology with advice from AS and IS. NWH made significant contributions to developing the multiwell MEA data analysis pipeline. NC analysed the high-density multielectrode array data and wrote parts of the methods section “Data analysis”. VF carried out all other experimental work and wrote the rest of the manuscript, with input from NWH, NC, AS and IS.

## Supplementary material

**Supplementary figure 1.**
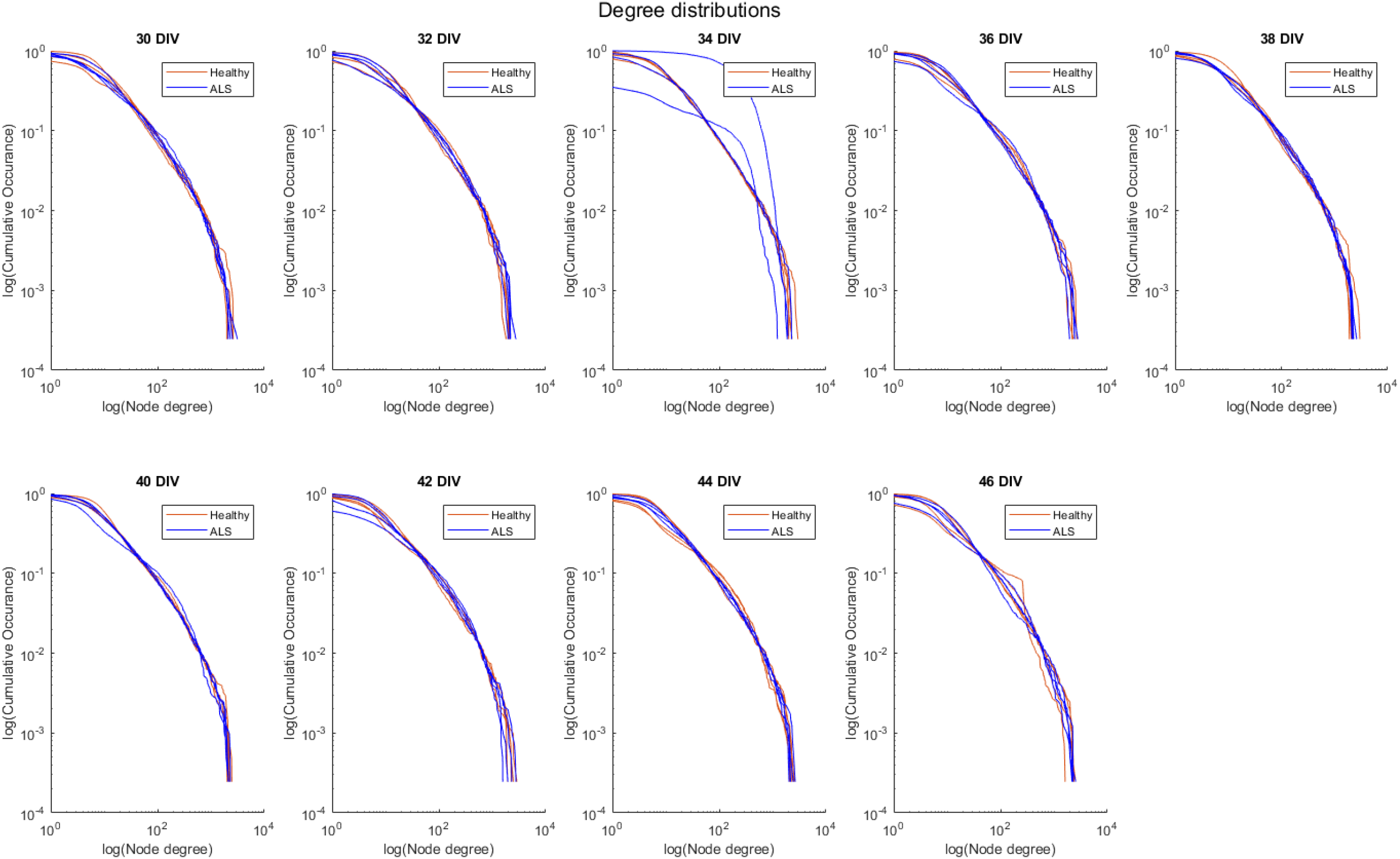
Motor neuron network degree distributions. Motor neuron networks on HD-MEAs have node degree distributions which appear to follow a power-law, i.e. linear in a log-log scale.

